# Diminished distractor filtering with increased perceptual load and sustained effort explains attention deficit in post-stroke fatigue

**DOI:** 10.1101/2022.03.17.484709

**Authors:** Annapoorna Kuppuswamy, Anthony Harris, William De Doncker, Adrian Alexander, Nilli Lavie

**Affiliations:** Department of Clinical and Movement Neuroscience, Institute of Neurology, UCL, United Kingdom; Institute of Cognitive Neuroscience, UCL, United Kingdom; Queensland Brain Institute, The University of Queensland, Australia; San Fernando General Hospital, South West Regional Health Authority, Trinidad and Tobago

**Keywords:** post-stroke fatigue, visual perception, distractor, attention, perceptual load

## Abstract

**Objectives:** Post-stroke fatigue (PSF) is a prevalent symptom associated with attention deficits. However, it is currently unclear what drives these. Here we applied Load Theory of Attention to investigate the role of perceptual load in the relationship between attention, distraction, and fatigue levels in PSF.

**Methods:** Thirty-two chronic stroke survivors performed a selective attention task under different perceptual load levels. Neural responses to targets and distractors were measured with frequency-tagged EEG responses.

**Results:** We showed that Fatigue Severity Scale-7 (FSS-7) scores are predictive of slower responses, irrespective of the task level of perceptual load. Although average distractor frequency (10Hz) power was reduced with higher perceptual load, the reduction in 10Hz power with increasing task load, was smaller in those with higher FSS-7 scores. Higher FSS-7 scores were also associated with increased 10Hz distractor response as time progressed, both within task-trials and across the experiment duration.

**Conclusion:** The results are consistent with previous reports of slower processing speed and sustained selective attention deficits. Importantly they demonstrate that higher fatigue severity is associated with a reduced ability to suppress neural response to irrelevant distractors stimuli when the task demands more attention due to its increased perceptual load. An account for attention deficit in PSF based on a distractor filtering deficit that emerges with increased perceptual load and sustained effort can explain a myriad of PSF symptoms (e.g. sensory perceptual overload, and difficulties to concentrate).

**Key messages:**

- **What is already known on this topic**
  ➢ Post-stroke fatigue involves mental fatigue which is exacerbated in cognitively demanding task, as well as difficulties in concentrating and sustaining attention on a task.
  ➢ However previous research has not converged on a clear account of the nature of attention deficits in post-stroke fatigue, as well as on how these relate to the level of perceptual demand in a cognitive task
- **What this study adds**
  ➢ The study suggests that attention deficits in post-stroke fatigue are attributed to a failure to attenuate neural responses to distractors with increased perceptual load in the task, as well as with increased demands on sustained attention with a longer task engagement
  ➢ This pattern points to a distractor filtering deficit that is magnified by perceptual load and sustained mental effort as main drivers of attention deficits in post-stroke fatigue **How this study might affect research, practice or policy**
  ➢ This study enhances our understanding of attentional deficit in post-stroke fatigue, within an integrative account that explains mental fatigue symptomology

## Introduction

Post-stroke fatigue (PSF) is a prevalent and persistent symptom after stroke characterized by severe debilitating exhaustion, both physically and mentally^1,2^. People with high levels of PSF report increased fatigue when performing cognitively demanding activities^3^ as well as problems concentrating^4^, often complain that they are unable to filter out irrelevant background stimuli. For example, they cannot carry a conversation in a crowded environment (e.g. a party), and find visual clutter overwhelming, resulting in avoidance of public spaces^1,5,6^. These are suggestive of an attention deficit, however previous studies led to mixed results. Whereas some studies clearly indicated deficits in the ability to sustain attention over time^4,7^, as well as in selective attention and ‘executive control’ over irrelevant distraction^3,4^, others reported generalized cognitive impairment expressed in slower processing speed and some memory deficits, but no impairment in selective attention and executive control^8–11^.

Here we applied Load Theory framework of selective attention and cognitive control^12,13^ to clarify the nature of attention deficit in PSF. In this model, due to limited capacity both for perception and cognitive executive control, perceptual processing proceeds on all stimuli including distractors, until capacity runs out. Perceptual load in the task is therefore a critical determinant of the ability to pay focused selective attention in healthy adults. Irrelevant distractors cannot be ignored and elicit neural response and behavioral interference in tasks of low perceptual load, but their processing is diminished with increased perceptual load that exhausts all perceptual capacity^12,14,15^. Cerebral metabolism limits on neural computations have been shown as the underlying limited resources explaining limited perceptual capacity^16^. Drawing on PSF research, indicating limited energy^17^, and increased mental fatigue developing in the course of performance in cognitive demanding tasks^3^ as well the complaints of PSF sufferers about over-stimulation, we hypothesized that perceptual load may be a critical factor in their attention deficits.

We set thus out to investigate interactions between attention, distraction, perceptual load and fatigue in chronic stroke using a selective attention task with frequency tagged EEG responses to targets and distractors^18^.

## Methods

Following written informed consent in accordance with the Declaration of Helsinki, thirty-two stroke survivors (Table 1) participated in this cross-sectional observational study approved by the London Bromley Research Ethics Committee (REC reference number: 16/LO/0714). Inclusion criteria: (1) first-time ischemic or hemorrhagic stroke (2) stroke >3 months at time of testing. Exclusion criteria (3) other neurological disorder (4) on anti-depressants or centrally acting medication (5) clinically depressed or Hospital Anxiety and Depression Scale (HADS) scores ≥11 (6) sensory impairment and neglect (7) hand grip strength and nine-hole peg test (NHPT) of the affected hand <60% of unaffected hand [Table 2].

**Table 1:** Clinical characteristics of the stroke across all study participants taken from the discharge summary hospital notes of each individual stroke survivor. Lesion location and hemisphere affected was defined based on the vascular territory affected by the stroke. The mean and standard deviation of the time post-stroke, taken in years, from the time of the stroke to the time of participation in the study, indicating that the studied cohort were all chronic stroke survivors.

|  |  |
| --- | --- |
| **Stroke survivors total** | 32 |
| **Hemisphere Affected** |  |
| Right | 13 (40.62%) |
| Left | 19 (59.38%) |
| **Stroke Type** |  |
| Ischemic | 29 (90.62%) |
| Hemorrhagic | 3 (9.38%) |
| **Vascular Territory Affected** |  |
| ACA | 0 (0.00%) |
| MCA | 23 (71.88%) |
| PCA | 2 (6.25%) |
| Brainstem/Cerebellum | 7 (21.88%) |
| **Time Post-Stroke (years)** |  |
| Mean (SD) | 4.60 (3.52) |
| ACA = Anterior Cerebral Artery; MCA = Middle Cerebral Artery; PCA = Posterior Cerebral Artery |  |

**Table 2:**
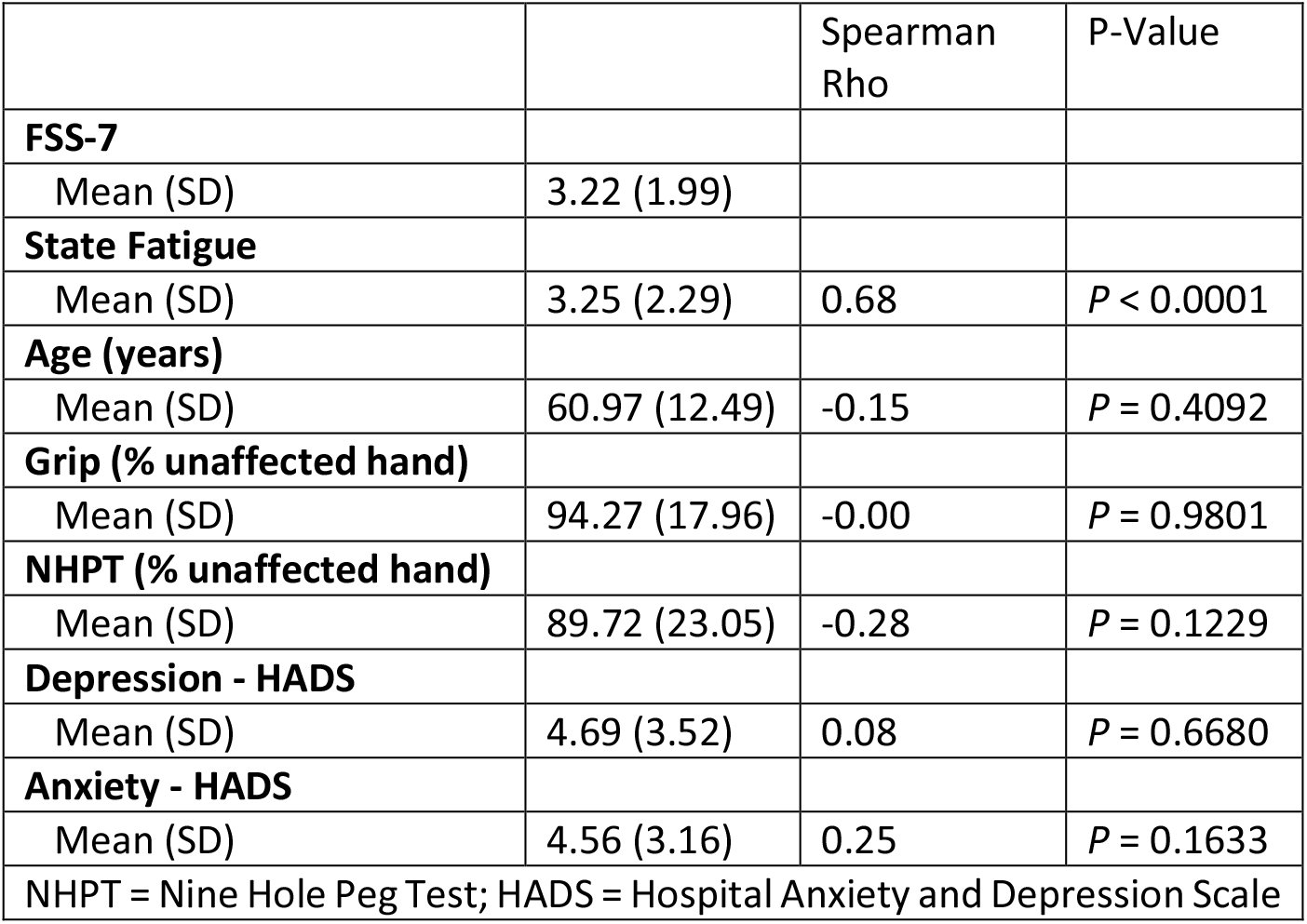
Demographics of the study participants indicating the mean and standard deviation of the various fatigue, motor, cognitive and mood scores. Spearman correlations were used to assess the association between FSS-7 and all continuous measures. The spearman rho and associated p-values indicate the relationships between FSS-7 score and the respective measures. There is a significant positive association between trait (FSS-7) and state fatigue. Non-significant relationship between FSS-7 and all measures indicate that the group studied, suffered from what is known as ‘primary’ fatigue i.e. that which is not secondary to obvious motor or mood impairments. Mean grip and NHPT scores of approximately 90% reflects the minimal impairment of the group.

|  |  | Spearman Rho | P-Value |
| --- | --- | --- | --- |
| **FSS-7** |  |  |  |
| Mean (SD) | 3.22 (1.99) |  |  |
| **State Fatigue** |  |  |  |
| Mean (SD) | 3.25 (2.29) | 0.68 | $P < 0.0001$ |
| **Age (years)** |  |  |  |
| Mean (SD) | 60.97 (12.49) | -0.15 | $P = 0.4092$ |
| **Grip (% unaffected hand)** |  |  |  |
| Mean (SD) | 94.27 (17.96) | -0.00 | $P = 0.9801$ |
| **NHPT (% unaffected hand)** |  |  |  |
| Mean (SD) | 89.72 (23.05) | -0.28 | $P = 0.1229$ |
| **Depression - HADS** |  |  |  |
| Mean (SD) | 4.69 (3.52) | 0.08 | $P = 0.6680$ |
| **Anxiety - HADS** |  |  |  |
| Mean (SD) | 4.56 (3.16) | 0.25 | $P = 0.1633$ |
| NHPT = Nine Hole Peg Test; HADS = Hospital Anxiety and Depression Scale |  |  |  |

### Fatigue

Trait fatigue was measured using Fatigue Severity Scale -7 (FSS-7)^19^ and state fatigue was measured using visual analogue scale (VAS, 0-10).

### Selective attention task stimuli

Visual stimuli were presented on a Dell U240 24-inch monitor, at a screen resolution of 1280×768, and a refresh rate of 60 Hz. Participants were seated 70 centimeters from the monitor and made their responses using a standard USB keyboard. The experiment was controlled using the Psychophysics Toolbox for Matlab^20,21^, running on a Windows computer.

The experiment was a full crossing of three perceptual load conditions (Low, Medium, and High load) with two peripheral stimulation conditions (No flicker vs. Flicker). In all conditions, participants were presented with streams of 32 coloured crosses (2.5° tall, 1.5° wide, 0.3° thick). Each cross could appear in one of 6 colours and each was either upright or inverted (**Figure 1B**). The colours were as follows: red (RGB: 255, 0, 0), green (RGB: 0, 255, 0), blue (RGB: 0, 0, 255), brown (RGB: 156, 102, 31), yellow (RGB: 255, 255, 0), and purple (RGB: 160, 32, 240). For upright and inverted crosses, the horizontal bar was 0.4° above or below the midline, respectively. All stimuli were presented on a grey background (RGB: 127, 127, 127). Each cross was presented for 250ms, followed by a 500ms inter-stimulus interval. Each cross stimulus was presented on a central disk (2.5° x 2.5°), flickering between dark grey (RGB: 64, 64, 64) and light grey (RGB: 192, 192, 192) at a frequency of 4 Hz (one change every 250ms). In addition, on Flicker trials, the central stream of crosses was surrounded by a ring of unattended counterphasing visual checkerboards (**Figure 1A**). These checkerboards spanned from 2.5° to 14° into the periphery, divided into 19 concentric rings of 0.6° each, and were made up of 56 radial sections, each spanning 6.4° of radial angle. Successive cells of this checkboard were each alternately coloured light or dark grey, and their colours counterphased at a rate of 10 Hz. On No-flicker trials, this checkerboard was replaced by the grey background colour. Such checkerboards elicit strong neural responses across a range of visual areas^15^ at the precise frequency of the visual flicker^18,22^.

**Figure 1.**
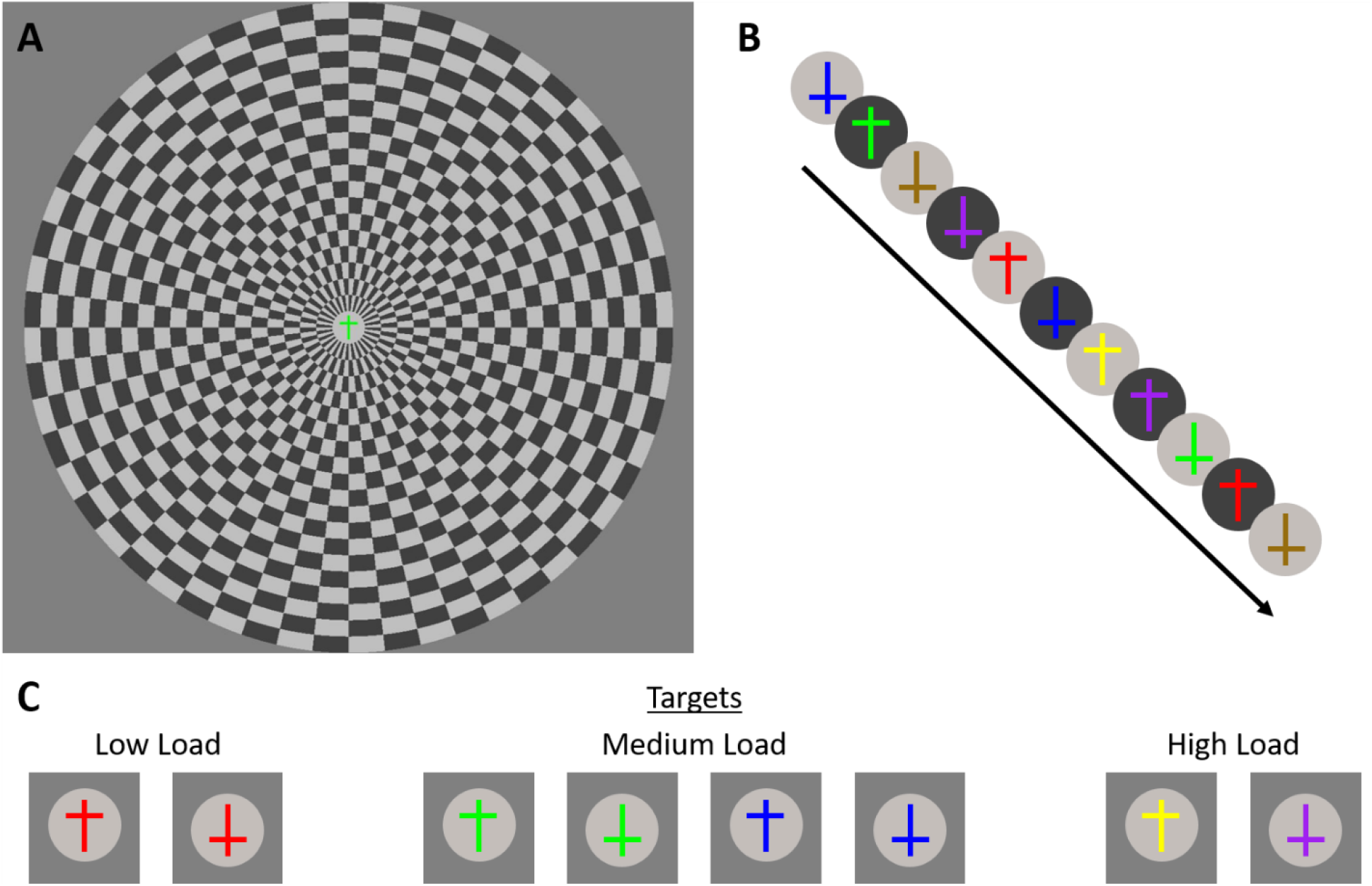
Stimuli: A) Participants attended a central stream of crosses, while, in the Flicker condition, they ignored a peripheral flickering checkerboard. This checkerboard was absent in the No flicker condition. B) The central stream contained upright and inverted crosses that varied among six different colours. These changed every 750ms. C) In the low load condition, participants were to respond whenever they saw a red stimulus. In the medium load condition, they were looking for any green or blue stimuli. Finally, in the high load condition, targets were upright yellow crosses and inverted purple crosses. The colour of the disk behind the cross was not target relevant.

### Procedure

Participants fixated on a fixation marker, and when the trial stream appeared, and a target was detected, participants responded by pressing the spacebar. Participants were required to withhold responses to all nontarget stimuli. Each stream contained 4 targets and 28 nontargets (trial duration=24 seconds). The position of targets in the stream was restricted such that the first 2 items could not be targets, nor could the last item, and no two targets could appear back-to-back.

Three levels of perceptual load were included (**Figure 1C**). In the low load condition, the targets were upright and inverted red crosses, in the medium load condition, targets were any (upright or inverted) green or blue cross and in the high load condition, targets were upright yellow crosses and inverted purple crosses.

Participants completed three practice trials (one for each load level), then 48 trials of the main task. The experiment was composed of 8 trials from each of the 6 different conditions (3 load conditions crossed with 2 flicker conditions). Each trial contained 4 targets, giving 32 targets per condition. Participants received a self-paced break after every 10 trials of the main task.

### EEG Recording

Whole-scalp electroencephalography (EEG) data was recorded using a 64-channel cap array (ActiCap, Herrsching, Germany) in accordance with the 10-20 international EEG electrode array and a BrainAmp EEG amplifier system (BrainProducts, Gilching, Germany). During online recordings, channels FCz and AFz were used as the reference and ground respectively. Vertical and horizontal electrooculogram (EOG) recordings were used to capture horizontal eye movements and blinks. The EEG signal was sampled at 1 kHz and event markers were sent from the stimulus presentation PC to the BrainAmp amplifier via the TriggerBox with millisecond precision.

### EEG Analysis

Using a combination of EEGLAB^23^ and custom Matlab scripts, data were imported, and bad channels were identified using EEGLAB’s default kurtosis-based automated procedure. Any bad channels were then interpolated using spherical spline interpolation followed by re-referencing against the grand average of all scalp electrodes. Epochs were extracted from 2 seconds before the beginning of each trial until the end of each trial, and baseline corrected against the 2 seconds before trial onset. Steady-state visual evoked potential (SSVEP, a.k.a., ‘frequency tag’) amplitudes were calculated across the trial period by means of a Fast Fourier Transform on each 1-second period of trial data (24 per trial) which were averaged within the trial, then across trials within each condition. To increase the signal-to-noise ratio of our data, results were averaged over a cluster of posterior electrodes (electrodes Iz, O1/2, Oz, and POz). The SSVEP responses to the background flicker (10 Hz frequency) as well as the frequency of flicker at the target location (4 Hz) were used for statistical analysis.

### Statistical Analysis

Spearman rank correlations were used to identify associations between FSS-7 and state fatigue, age, grip strength, NHPT, HADS–Depression, HADS–Anxiety and Time Post-Stroke. Wilcoxon rank sum tests were used to identify the association between FSS-7 score and categorical measures of sex, hemisphere affected, type of stroke and vascular territory affected.

Two-way repeated measures ANOVAs with the factors of flicker (On/Off) and Load (Low, Medium, High) were conducted on a) response time, b) accuracy, c) 4 Hz power (target frequency), and d) 10 Hz power (distractor frequency). Greenhouse-Geisser epsilon adjustment corrected for deviations from sphericity. Post-hoc pairwise t-tests with Bonferroni corrections identified main effects and interactions.

Multiple linear regressions were used to investigate the effect of a) FSS-7 and load (Low, Medium, High) on response time and accuracy b) FSS-7 on load-related brain responses, operationalized as the change in 10Hz power between the load levels (Δ10Hz = 10Hz medium - 10Hz low, 10Hz high – 10 Hz low) across flicker (On/Off) conditions. The best fitting model was determined using the Bayesian Information Criterion (BIC), with a lower BIC indicating a better fitting model. Assumptions of normality and homoscedasticity of the residuals for each linear regression model were assessed visually using quantile-quantile normal plots and fitted-versus residual-value plots.

To measure the effect of FSS-7 on 10Hz power over time, both within a trial, and over the course of the experiment, a linear mixed effects analysis was performed individually on each flicker condition (On and Off). The models were specified with a random effect for participants, and fixed effects of load, time within trial (early vs late; 2 × 12 sec bins), trial number, and FSS-7, as well all two and three-way interactions of time within trial, trial number, FSS-7.

## Results

### Clinical Characteristics of Stroke

There was no difference in FSS-7 score between right [median=4.5 (IQR = 4.2)] and left hemisphere [median=1.9 (IQR=3.0)] strokes (p=0.4, effect size r=0.14), ischaemic [median=1.9 (IQR=3.7)] and hemorrhagic [median=2.8 (IQR=2.0)] strokes (p=1, effect size r=0.01), and MCA [median=2.8 (IQR=4)], PCA [median=3.8 (IQR=1.9)] and brainstem/cerebellar [median=1.9 (IQR=2.6)] strokes (p=0.6, effect size ɳ^2^=- 0.04). there was also no correlation between FSS-7 and time post stroke (spearman ρ=0.25, p=0.17).

### Response Time

There was a main effect of load (F_(1.61,50.04)_=174.00, p<0.001, ɳ^2^=0.43), no effect of flicker (F_(1,31)_=0.49, p=0.49, ɳ^2^<0.001) and no interaction between load and flicker (F_(1.58,49.11)_=0.40, p=0.63, ɳ^2^<0.001). Post-hoc tests revealed a significant increase from low to medium (t(31)=-11.00, p<0.001, d=-1.94), and from medium to high (t(31)=-9.67, p<0.001, d=-1.71), as well as from Low to High (t(31)=-16.70, p<0.001, d=- 2.96) [Fig 2A].

**Figure 2.**
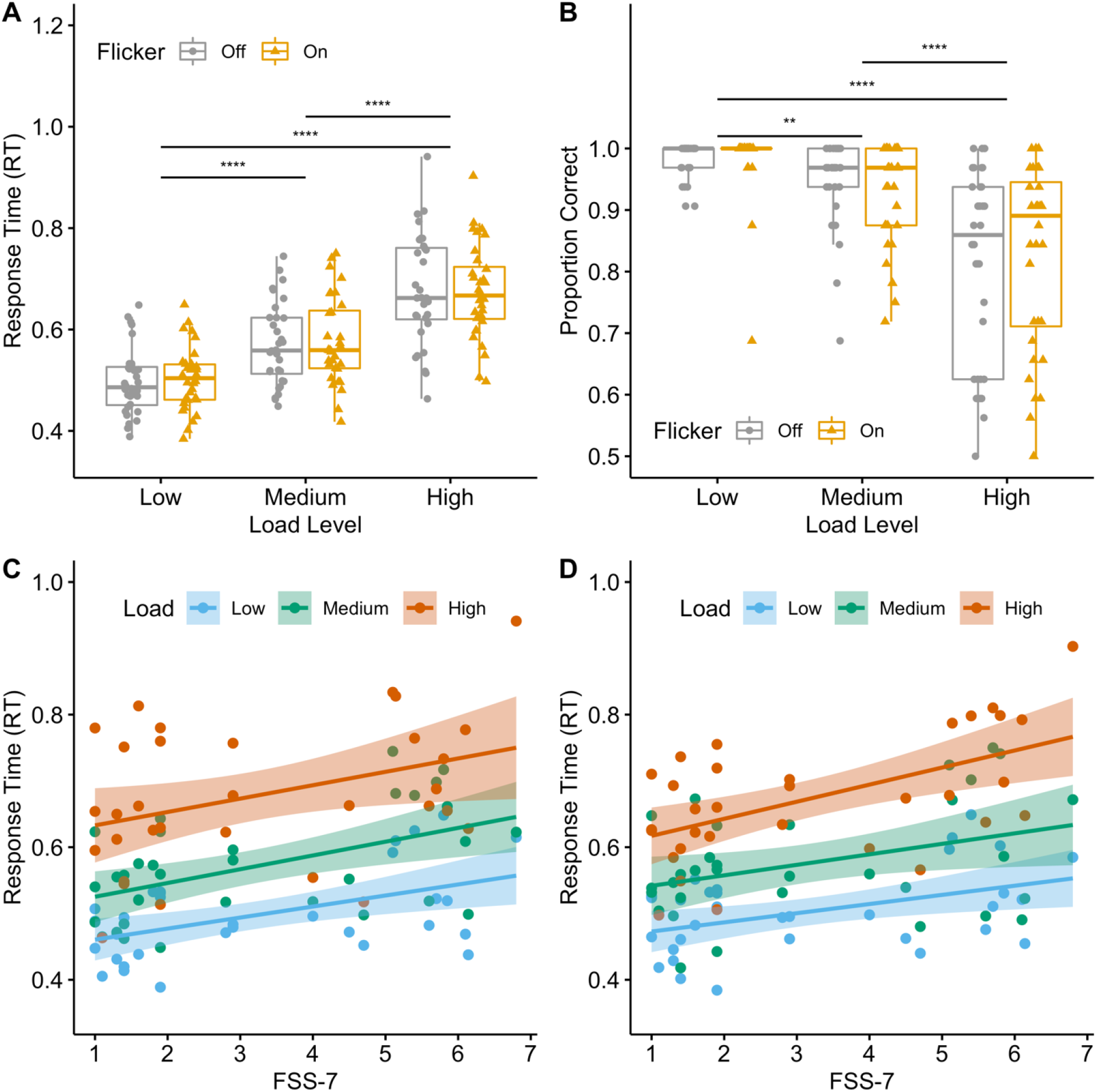
Response Time & accuracy: In **2A**, the time taken to respond to a target stimulus (Response Time – y-axis) is shown for all three load conditions (x-axis) in the presence (ochre) or absence (grey) of a flickering distractor. RT was significantly longer as load increased, with no difference across flicker conditions. In **2B**, the response accuracy (proportion correct – y-axis) is shown for all three load conditions (x-axis) in the presence (ochre) or absence (grey) of a flickering distractor. Proportion correct was significantly lower with increasing load, with no effect of flicker. In **2C**, RT (y-axis) is shown to increase significantly with increase in fatigue levels (FSS-7, x-axis) in the flicker Off condition. In **2D**, RT (y-axis) is shown to increase significantly with increase in fatigue levels (FSS-7, x-axis) in the flicker On condition.

The linear regression model with response time as dependent variable and FSS-7, state fatigue and load (Low, Medium, High) as predictors was significant (F_(4,91)_=33.92, p<0.001) with an adjusted R^2^ of 0.58 and significant main effects of trait fatigue (F_(1,91)_=29.22, p<0.001, ɳ^2^=0.13), state fatigue (F_(1,91)_=7.11, p=0.01, ɳ^2^=0.03) and load (F_(2,91)_=50.90, p<0.001, ɳ^2^=0.44). Beta coefficients for significant predictors were, FSS-7 (β=0.03, p<0.001, CI[0.02,0.04]), state fatigue (β=-0.01, p=0.01, CI[-0.02,0.00]) and Load (β_Medium-Low_=0.07, p<0.001, CI[0.04,0.11]; β_High-Low_=0.18, p<0.001, CI[0.14,0.21]).

### Accuracy

There was a main effect of load (F_(1.41,43.73)_=33.95, p<0.001, ɳ^2^=0.32), no effect of flicker (F_(1,31)_=0.25, p=0.62, ɳ^2^<0.001) and no interaction between load and flicker (F_(2,62)_=1.35, p=0.27, ɳ^2^=0.005). Post-hoc tests show a significant decrease from low to medium (t(31)=3.94, p=0.001, d=0.70), from medium to high (t(31)=5.20, p<0.001, d=0.92), and from low to high (t(31)=6.64, p<0.001, d=1.17).

The linear regression model with accuracy as the dependent variable and FSS-7, state fatigue and load (Low, Medium, High) as predictors, was significant (F_(4,91)_=12.60, p<0.001) with an adjusted R^2^ of 0.33 and a main effect of load (F_(2,91)_=24.74, p<0.001, ɳ^2^=0.35) but no effect of trait fatigue (F_(1,91)_=0.06, p=0.80, ɳ^2^<0.1) or state fatigue (F_(1,91)_=0.61, p=0.44, ɳ^2^<0.1). Load was a significant predictor of accuracy (β_Medium-Low_=-0.05, p=0.04, CI[-0.10,-0.01]; β_High-Low_=-0.16, p<0.001, CI[-0.21,-0.12])with no effect of FSS-7 (β=- 0.002, p=0.80, CI[-0.02,0.01]) or state fatigue (β=0.005, p=0.44, CI[-0.01,0.02]).

### 4 Hz Power (Target Frequency)

There was no significant effect of load (F_(1.6,50.6)_=0.49, p=0.58, ɳ^2^=0. 0003, figure 4A), or flicker on 4Hz power (F_(1,31)_=1.06, p=0.31, ɳ^2^=0.0007, figure 4A) and no interaction between load and flicker (F_(2,62)_=0.90, p=0.41, ɳ^2^=0.0005).

### 10 Hz Power (Distractor Frequency)

There was a main effect of load (F_(2,62)_=12.64, p<0.001, ɳ^2^=0.20, figure 3A, 4B and 4C) and flicker (F_(1,31)_=28.64, p<0.001, ɳ^2^=0.28, figure 3B and 4B) but no interaction between load and flicker (F_(2,62)_=0.17, p=0.841, ɳ^2^<0.001). Post-hoc tests revealed a significant decrease from Low to High Load (t(31)=4.25, p<0.001, d=0.75), and Medium to High Load (t(31)=3.32, p=0.007, d = 0.59), but no decrease from Low to Medium Load (t(31)=2.01, p=0.16, d=0.36).

**Figure 3.**
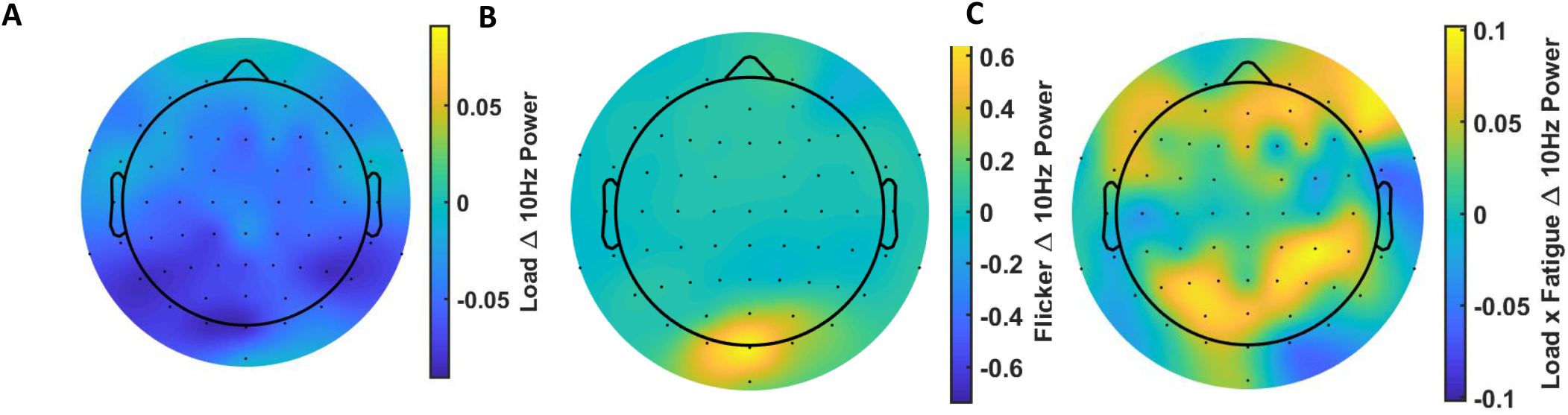
10hz power visualisation: Figure **3A** shows the change in 10Hz power between high and low load conditions. Figure **3B** shows the difference in 10Hz power between ‘Flicker’ and ‘No flicker’ conditions. Figure **3C** shows the difference between high (FSS-7>4) and low fatigue (FSS-7<4) in the reduction in 10Hz power between high and low load conditions.

**Figure 4.**
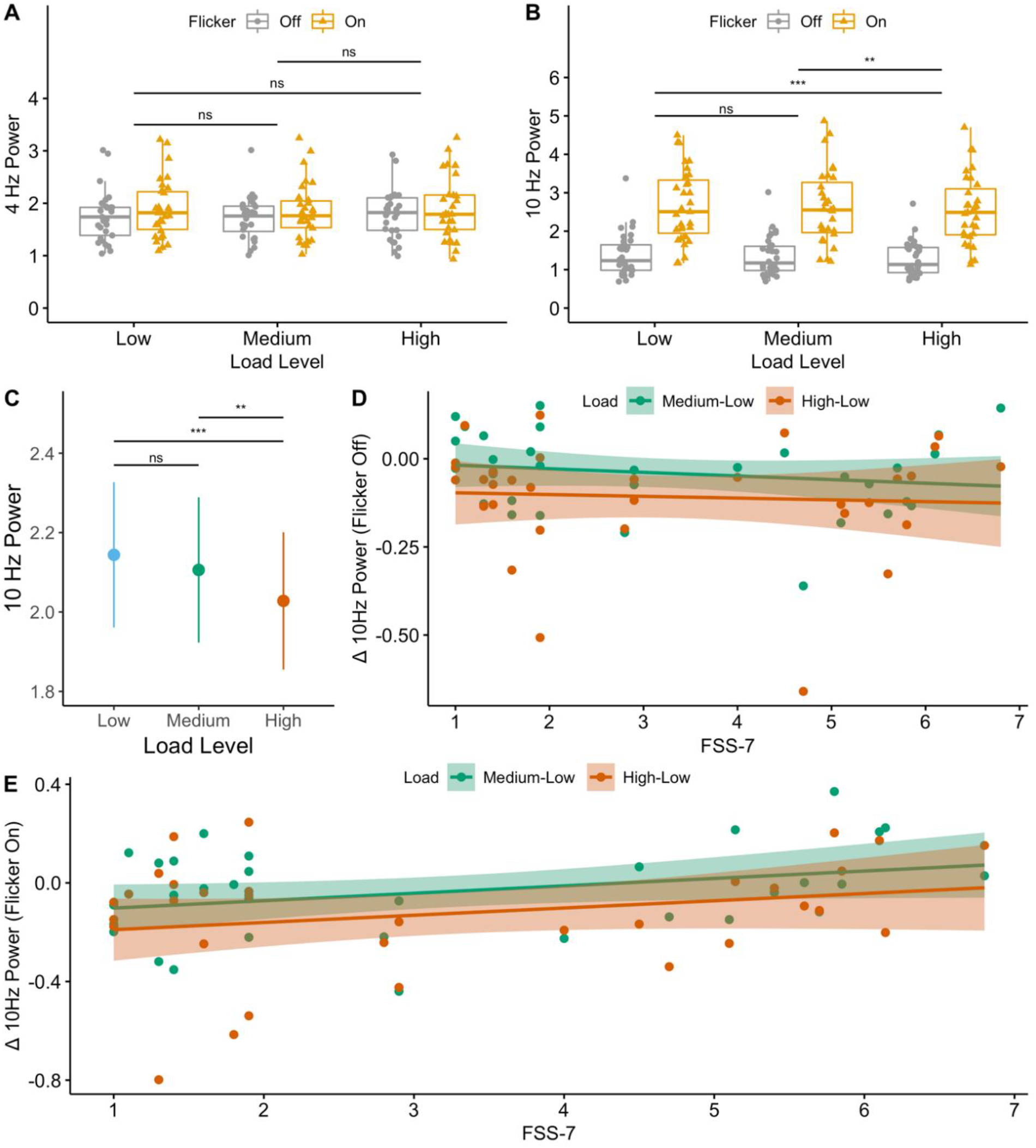
10Hz power: Figure **4A** shows that power at 4Hz is not significantly different between the two flicker conditions or between the load conditions. **4B** shows that power at 10Hz is significantly higher in the ‘flicker on’ condition when compared to ‘flicker off’ condition confirming the response to the distractor frequency. **4C** highlights the significant reduction in 10Hz power as the task load increases. In Fig **4D** and **4E**, the decrease in 10Hz power (Δ10Hz power) from medium to low (green) and high to low (red) load is correlated to fatigue levels (FSS-7) for the flicker Off and flicker On conditions respectively. In the flicker On condition, the higher the fatigue, the smaller is the decrease in distractor related power as load increases whether from low to medium or from low to high. This was not observed in the flicker Off condition.

The multiple linear regression model for the Flicker Off condition was not significant (F_(3,60)_=2.27, p=0.09) with an adjusted R^2^ of 0.06 (figure 4D) and no main effect of FSS-7 (F_(1,60)_=0.36, p=0.55, ɳ^2^=0.005), load (F_(1,60)_=3.70, p=0.06, ɳ^2^=0.06), or state fatigue (F_(1,610)_=2.37, p=0.13, ɳ^2^=0.04).

For the Flicker On condition, the multiple linear regression model was significant (F(3,60)=3.55, p=0.02) with an adjusted R^2^ of 0.11 (Figure 4E) and a main effect of trait fatigue (F_(1,60)_=6.89, p=0.01, ɳ^2^=0.09) but no effect of load (F_(1,60)_=3.15, p=0.08, ɳ^2^=0.04) and no effect of state fatigue (F_(1,60)_ = 2.15=6, p=0.15, ɳ^2^=0.03. FSS-7 was a significant predictor of Δ10Hzpower (β=0.05, p=0.01, CI[0.01,0.09], figure 3C) while load (β=-0.09, p=0.08, CI[-0.19,0.01]) and state fatigue (β=-0.03, p=0.15, CI[-0.06,0.01]) were not significant predictors of Δ10Hz Power.

### Effect of time on 10Hz power

In the flicker off condition, there was a significant effect of the intercept (β=.37, *p*<0.001), indicating 10 Hz power differed from zero as expected, but no other significant effects (all *p*s>0.335). In contrast, for the flicker on condition, we found significant effects of the intercept (β=0.88, *p*<0.001) and load (β=-0.03, *p*=0.006), indicating that, as load increased, 10Hz power decreased. We also found significant interactions between fatigue and trial number (β=0.002, *p*=0.039) and between fatigue and time within trial (β=0.04, *p*=0.032) (Figure 5). None of the other effects were significant (all *p*s>0.252).

**Figure 5.**
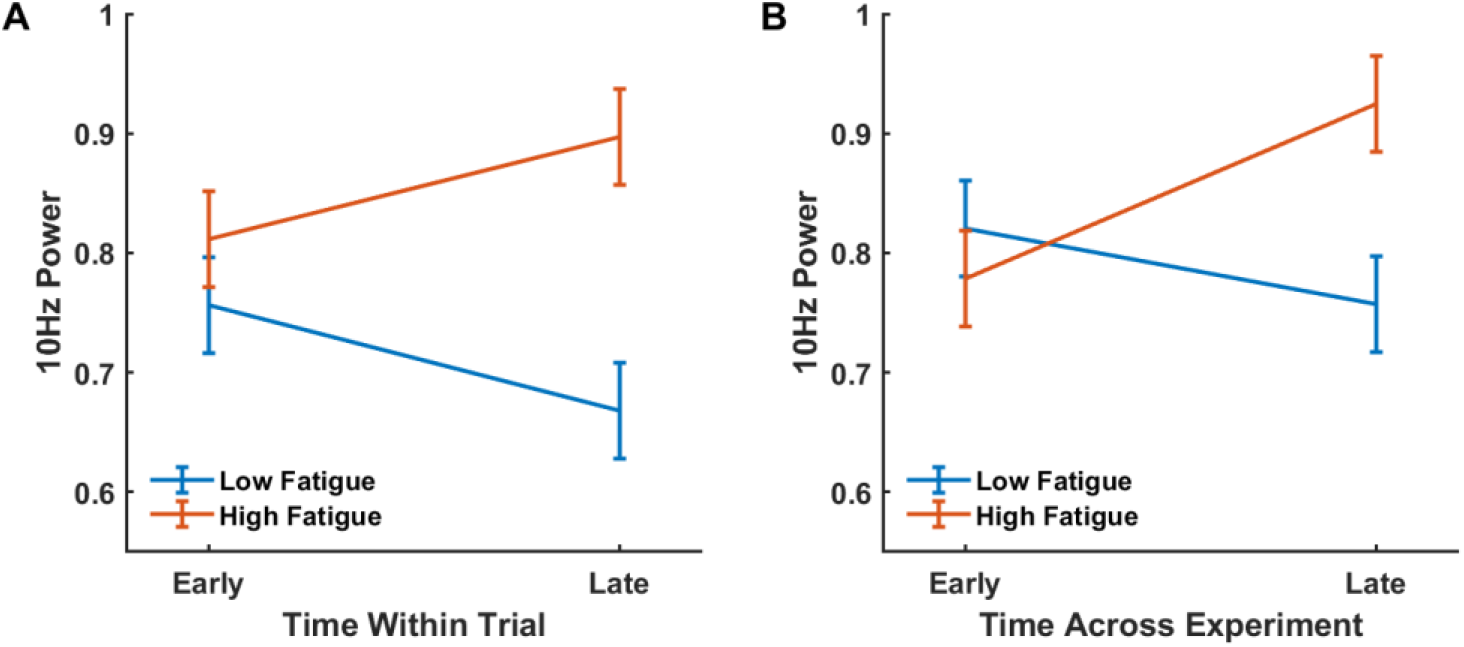
Interactions between Fatigue and Time in the Flicker On condition: Figure **5** shows that power at 10 Hz increases across time – both within a trial (**A**) and across trials (**B**) – for those with high fatigue, while it decreases across time for those with low fatigue. These results suggest those with low fatigue are able to exert greater attentional control as time goes on, while attentional control progressively worsens for those with low fatigue. Here, low and high fatigue are estimated 1SD below and above the mean fatigue score, respectively. Early and Late refer to the first and second half of a trial in **A** and the first and second half of the experiment in **B**.

## Discussion

In thirty-two minimally impaired, non-depressed, chronic stroke survivors with no visual neglect or extinction, we show that trait fatigue level is predictive of slower responses in a visual attention task, including when controlling for state fatigue. As perceptual load increased, there was a significant increase in response times too, however, there was no interaction between trait or state fatigue and the effect of load on response times. Accuracy of responses dropped with increasing load, but there was no effect of fatigue on accuracy. There was no effect of distractor on response times. However, EEG analysis revealed that 10 Hz power (the frequency of the flickering distractor) was significantly higher in the distractor ‘On’ condition, with increasing load diminishing 10 Hz power. The reduction in 10Hz power with increasing load was smaller in those with greater trait fatigue, with state fatigue not having any significant influence. We also show that as time progresses within a trial, and over the entire experiment, there is diminished suppression of distractor stimuli in those with greater trait fatigue. Similar to previous reports^18^ we did not observe any effect of load or distractor on 4 Hz power, the frequency at which the target stimulus was presented.

The finding that trait fatigue is associated with slower response speed is consistent with much previous research^4,7^. However, our novel finding of a reduction in the load effect on distractor processing (10Hz power in flicker On condition) in high trait fatigue is of most importance as it clarifies the relationship between perceptual load, distractor processing and fatigue. Together with the findings that the load modulation of the 10Hz power in the flicker Off (in which case, 10hz power can only be driven by endogenous oscillatory activity in the alpha range) was not associated with fatigue severity, these findings indicate that fatigue impairment in attention specifically implicates a reduced ability to attenuate distractor processing as the task processing becomes more demanding on perceptual processing, with increased perceptual load. This conclusion can explain patients’ reports of finding it hard to concentrate^3^, and of being overwhelmed by a visual scene that consists of several stimuli^1,5^. It may also suggest an account for previous discrepancies in research, by pointing to the critical factor of perceptual load in attention deficits associated with fatigue.

Our findings that the flickering distractor induced a neural response in the absence of behavioural interference effects also points to the importance of assessing neural responses to distracting stimuli as a clearer measure of any alterations in attention due to fatigue. Moreover, the findings that fatigue is associated with a failure to attenuate neural response to a non-interfering distractor (as measured with task performance), may provide further support to recent proposals that a fundamental deficit that explains both cognitive and motor aspects of post-stroke fatigue, is an alteration in perception - in vision, audition and proprioception^24,25^. This proposal suggests that incoming sensory information that is normally attenuated, (whether self-generated or external task-irrelevant stimuli), is poorly attenuated in those with high fatigue. For example, poor attenuation of proprioceptive input from contracting muscles can lead to a muscle contraction being experienced as effortful. Similarly, inability to attenuate processing of task-irrelevant visual stimuli can make a visual task feel more effortful. Despite being effortful, our previous studies show that there is no significant effect on performance metrics such as reaction times^26^ and completion of motor tasks^27^ suggesting a dissociation between measures of performance and reported effort, such that although the patients with chronic fatigue are able to achieve the same levels of performance these entail more effort for them. In the current study, lack of modulation of distractor effects by fatigue on behavioural measures, but increased neural response to the distractor with increased severity of fatigue is in further support of the dissociation between perceived effort and performance. Moreover, the findings that a non-interfering distractor shows a stronger brain response even under higher load conditions, suggests that in situations where distractors carry greater meaningfulness and behavioural relevance (such as a crowded shopping center) such perceptual intrusions may be even stronger.

Our findings that greater fatigue severity was also associated with an increase in the 10 hz response to the distractor as the demands on sustained attention increase over the a trial duration as well as over the duration of the experiment is consistent with previous research pointing to deficits in sustained attention^4,7^. Moreover, since this effect was only found in the 10 hz response to a distractor (in the distractor On condition) it indicates that patients with greater fatigue exhibited a larger decline in executive network efficiency with greater demand on sustained attention than patients reporting lower levels of fatigue. Therefore, the ability to sustain attention, both for short (within a trial) and long periods of time (entire experiment), is compromised in those with high fatigue, specifically in the presence of task unrelated visual distractors. This further explains patient reports of being unable to perform activities in crowded environments.

As mentioned earlier our finding that trait fatigue is predictive of slowed response times is consistent with previous research. This effect may indicate either slower speed of perceptual processing of the task stimuli or a deficit associated with the greater sensation of motoric effort. Our additional findings that perceptual load affects response times irrespective of fatigue severity, is in support of the latter interpretation because higher perceptual load increases the demand on perceptual processing (specifically here requiring a more complex discrimination of visual feature combinations) and should thus have resulted in magnification of the effect on response speed if it reflected any effect of fatigue on perceptual processing. Mixed results in clinical studies of attention where only a single load level is tested may thus be explained by other driving factors, such as greater movement related perceived effort^27^, or a general drop in arousal levels which has previously been implicated in other classic post-stroke visual perceptual deficits such as hemi-spatial neglect^28^. The finding of state fatigue predicting faster response times after controlling for trait fatigue should be interpreted with caution. When measuring the effects of trait fatigue on response times, it is critical to control for state fatigue, but not vice-versa. Given the strong effect size of trait fatigue on response times, and the effect of state fatigue being in the opposite direction, suggests that trait and state fatigue are distinct measures and influence response times independently.

In summary, we show that attention impairment in post-stroke fatigue is attributed to a reduced attenuation of distractor processing with increased perceptual load, as well as increased distractor processing with increased demand on sustained attention in PSF, as time goes on, throughout the experiment and even throughout a trial. Finally, our finding of slowing in response speed associated with greater fatigue severity, together with the equivalent perceptual load effect on responses across PSF symptom severity may be attributed to motor related perceived effort, rather than slower perceptual processing. Overall our findings clarify the nature of attention impairment in PSF, and can also explain the experience of cognitive fatigue, and difficulties to concentrate together with feeling that a stimulating environment might be overwhelming for the large group of PSF sufferers.

